# A Physics-Constrained Marcus-Theory Optimization Framework Restores Parameter Identifiability in Deep-Pocket Oncogenic Drug Kinetics: A KRAS G12C and STAT3 Case Study

**DOI:** 10.64898/2026.10.28.762299

**Authors:** L. M. Cluchagues Abam, I. H. Mfabo Kameni, F. Eya’ane Meva

## Abstract

**Background:** More than 90% of drug candidates fail preclinically, partly because empirical dose–response models such as the Hill equation ignore microenvironmental effects inside the deep binding pockets of targets such as mutant KRAS G12C and STAT3. Bulk water is hypothesized to raise the reorganization energy (*λ*_ext_).

**Objective:** To develop a physics-constrained framework, the *Cluchagues-Marcus Framework*, that links local dielectric permittivity to ligand–pocket kinetics, and to determine which constraints restore parameter identifiability.

**Methods:** Non-adiabatic Marcus kinetics were coupled to a Hill-type equation derived from a steady-state scheme, and water exclusion was modeled as a collapse of *ε*_local_ from 80.4 to 2.0. Pocket radii and published reference constants were used for two pockets (KRAS G12C, PDB 6UT0; STAT3 SH2, PDB 6NJS). Three propositions were proved and checked with synthetic data and a global sensitivity analysis, and the identifiability result was numerically stress-tested on real napabucasin viability curves from 701 cancer cell lines.

**Results:** Water exclusion lowered *λ*_ext_ by 90%. At Δ*G*° = −0.10 eV, *k*_C_ rose 8.0-fold (KRAS) and 3.6-fold (STAT3). Acceleration occurs if and only if 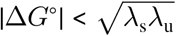. The model has exactly two non-identifiable parameter directions: fixing *v*_eff_ removes one and anchoring *K*_d_ removes the other. In real curves, fits were unchanged by the *K*_d_ anchor (SSE change < 10^−7^ %) while the fitted *k*_off,q_ varied by 3.5 log units, as predicted.

**Conclusions:** A dissociation-constant anchor makes the framework identifiable. The real-data stress test is not biological validation of the dielectric mechanism, which requires experiments.

## 1. Introduction

Hill–Langmuir dose–response models are the standard tool for estimating the potency of a ligand, but they describe equilibrium occupancy and contain no information about the kinetics of the transitions that occur once a ligand has entered a binding pocket [1,2]. Marcus theory, in contrast, gives transition rate constants that depend on the free energy of the step and on the solvent reorganization energy, which in a continuum description is controlled by the local dielectric permittivity [3–5]. Combining the two raises two mathematical questions. First, how does a change of the local dielectric constant change the rate constant, and under what conditions does it increase it? Second, when a Marcus rate constant enters a Hill-type equation, which of its parameters can be recovered from dose–response data [6,7]?

A motivating, but untested, scenario is a deeply buried binding pocket of an oncogenic target such as mutant KRAS (G12C), where bulk water (*ε*_local_ ≈ 80) could contribute a large reorganization energy [8–10]. Excluding water from such a pocket, for example with a hypothetical sub-nanometer dielectric shield of the size of atomically precise metal nanoclusters [11], would lower the local dielectric constant toward a hydrocarbon-like value of about 2. Green-synthesized metal nanostructures obtained with *Selaginella myosurus* extracts motivate this idea [12], but no sub-nanometer vector from this source has been demonstrated (Section 2.2). The present work does not test this scenario experimentally. It develops and analyses the resulting physics-constrained model, which we call the **Cluchagues-Marcus Framework**, and applies it to two deep-pocket targets, KRAS G12C and the STAT3 SH2 domain.

The contributions are as follows: (i) an exact condition on the binding free energy under which a reduction of the reorganization energy increases the Marcus rate constant, together with an upper bound on the rate constant (Proposition 1); (ii) a derivation of the kinetically corrected Hill equation from a steady-state scheme (Section 2.3); (iii) a proof that the model has exactly two structurally non-identifiable parameter directions, which are removed one by one when physical constants are fixed (Proposition 2); (iv) a numerical illustration and global sensitivity analysis; (v) two case studies (KRAS G12C and STAT3) showing how a dissociation-constant anchor and pocket geometry enter the framework; and (vi) a test of the identifiability result on real dose–response curves, with open code.

## 2. Methodology

### 2.1. Non-adiabatic Marcus kinetics

Transition kinetics within the dielectric-modified pocket were modeled with non-adiabatic Marcus theory [3, 5] (Eq. 1):

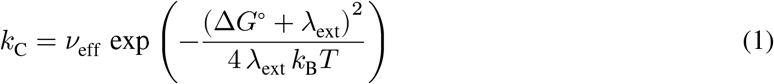

where *v*_eff_ is the effective attempt frequency (6.46 × 10^12^ s^−1^), which is also the upper bound of *k*_C_; Δ*G*° is the standard Gibbs free energy of the step, *k*_B_ is the Boltzmann constant and *T* is the absolute temperature (310.15 K) [13,14]. The model name denotes the present authors’ adaptation of Marcus’s electron-transfer theory; R. A. Marcus is not a contributor to this work.

### 2.2. Reorganization energy and the hypothetical dielectric shield

The external reorganization energy *λ*_ext_ as a function of the local dielectric permittivity *ε*_local_ was calculated from the continuum solvation expression based on Born’s model [4] (Eq. 2):

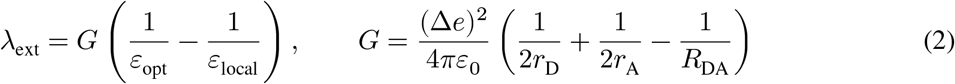

where *r*_D_ and *r*_A_ are the donor and acceptor radii, *R*_DA_ is the inter-particle distance, Δ*e* is the transferred charge and *ε*_opt_ is the optical dielectric constant (1.8) [4]. The geometric prefactor *G* was not assigned from structural coordinates; it was calibrated (*G* = 0.847 eV) so that *λ*_ext_ = 0.46 eV at *ε*_local_ = 80.4, the value adopted for the unshielded pocket. Because 1/*ε*_local_ increases when *ε*_local_ decreases, *λ*_ext_ is strictly smaller in the shielded pocket whenever the shielded dielectric constant is lower than the unshielded one (and larger than *ε*_opt_).

The dielectric shield was parameterized as a hypothetical sub-nanometer vector (*R* < 1.5 nm) occupying a deep binding cavity (*R*_pocket_ = 2.0 nm) [11, 15], and solvent exclusion was modeled as a step change from *ε*_local_ = 80.4 to *ε*_local_ = 2.0 [4, 9]. The cited *S. myosurus* study reported silver and silver chloride nanoparticles with crystallite sizes of 33.7 nm and 44.2 nm and a mean grain size of 58.81 nm by electron microscopy [12]; the sub-nanometer regime is therefore a hypothetical extrapolation to nanocluster dimensions [11], and no property of the plant extract is used as a model input.

#### Case-study pockets

Two ligand–pocket pairs were considered, each with the structure and the constant of the same ligand: adagrasib bound to K-Ras G12C in the switch-II pocket (PDB 6UT0, 1.94 Å resolution [16]; reference constant *K*_I_ = 1.43 µM, the inhibition constant measured by mass spectrometry on recombinant KRAS G12C [17,18]) and the degrader SD-36 bound to the STAT3 core (PDB 6NJS, 2.70 Å resolution [19]; reference constant *K*_i_ = 11 nM from a fluorescence-polarization binding assay [19]). Pocket radii were measured from the coordinate files with the script pocket_radius.py (provided with the code): the median distance from the ligand centroid to the protein heavy atoms within 4.5 Å of the ligand in chain A gave *R* = 0.72 nm for 6UT0 (ligand M1X; 86 lining atoms; maximum distance 1.22 nm) and *R* = 0.83 nm for 6NJS (ligand KQV; 71 lining atoms; maximum distance 1.65 nm). For a covalent inhibitor, *K*_I_ = (*k*_off_ + *k*_inact_) /*k*_on_ = *K*_d_ + *k*_inact_/*k*_on_, so *K*_I_ is an upper bound of the reversible encounter constant and not an equilibrium affinity; both constants are used only as reference scales for *K*_d_ in Eq. 3, and a reversible switch-I/II pocket inhibitor would be a more direct test. For a single-cavity Born model the prefactor *G* of Eq. 2 scales as 1/*R*; *G* was therefore calibrated at the KRAS reference pocket (*G* = 0.847 eV) and scaled by 0.72/*R* for STAT3 (*G* = 0.735 eV); because the calibration fixes the absolute scale, only the ratio of the two radii affects the results. This spherical single-cavity scaling is a simplification: real pockets are asymmetric and irregular, and geometric anisotropy would modulate *G* through shape factors that are left for future work. All other parameters were as in Table 2.

### 2.3. Kinetic scheme and dose–response equation

Eq. 3 follows from a minimal kinetic scheme [1, 20]. A ligand L binds the target P in a rapidly equilibrating step with dissociation constant *K*_d_, forming an encounter complex PL. PL is converted to an engaged state PL* with rate constant *k*_C_ (Eq. 1), and PL* is removed with rate constant *k*_off,q_ through a separate, non-reversing channel (relaxation or release of the engaged state). At steady state, d [PL^*^] /d*t* = *k*_C_[PL] – *k*_off,q_ [PL^*^] = 0, so [PL^*^] = (*k*_C_/*k*_off,q_) [PL] and the total ligand-bound target is [PL] (1 + *k*_C_/*k*_off,q_). The apparent steady-state constant is therefore *K*_app_ = *K*_d_/ (1 + *k*_C_/*k*_off,q_) = *K*_d_ *k*_off,q_/ (*k*_C_ + *k*_off,q_). The scheme describes a driven, non-equilibrium steady state (for example continuous cellular clearance or degradation of the engaged state), not a closed equilibrium, so Eq. 3 is a steady-state target-engagement model and *K*_d_-based expressions are apparent steady-state quantities. Taking the response (cellular depletion fraction, *Y*) to be proportional to total steady-state engagement and introducing a Hill coefficient *n* gives the modified Hill-type equation (Eq. 3). For *n* = 1 the factor applies exactly to *K*_d_; for *n* ≠ 1 it is applied to 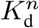, a phenomenological generalization [2, 21]:

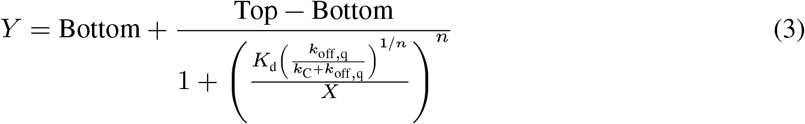

where *X* is the ligand concentration, *k*_off,q_ is the quantum-corrected off-rate constant, and Top and Bottom are the response asymptotes [2]. The effective half-maximal concentration is EC_50_ = *K*_d_ [*k*_off,q_/ (*k*_C_ + *k*_off,q_)]^1/*n*^.

### 2.4. Numerical illustration, identifiability analysis and sensitivity analysis

For each case study, synthetic dose–response data sets (*N* = 100 concentrations, log-spaced from 10^−10^ to 10^−4^ M) were generated from Eq. 3 with additive Gaussian noise (*σ* = 0.05), using the parameter values in Table 2. Data were fitted in Python with scipy.optimize.curve_fit (Trust Region Reflective algorithm, with bounds) [22] under three configurations: (i) six free parameters (Top, Bottom, *K*_d_, *n, k*_C_, *k*_off,q_); (ii) *v*_eff_ fixed, so that *k*_C_ follows from Eq. 1 (five free parameters); and (iii) *v*_eff_ and *K*_d_ fixed (four free parameters). Identifiability was assessed from the singular values of the Jacobian of the residuals at the optimum (central differences, parameters on a logarithmic scale where positive). The null dimension of a configuration was defined as the number of free parameters minus the number of singular values exceeding 10^−6^ times the largest [6,7]. The experiment was repeated for 200 independent noise realizations (seeds 0–199); a representative data set (seed 42) is shown in Figure 1. Goodness of fit was quantified by the coefficient of determination (*r*^2^). The propositions of Section 3.1 were also checked numerically (random parameter sets; Supplementary Code S1).

**Figure 1.**
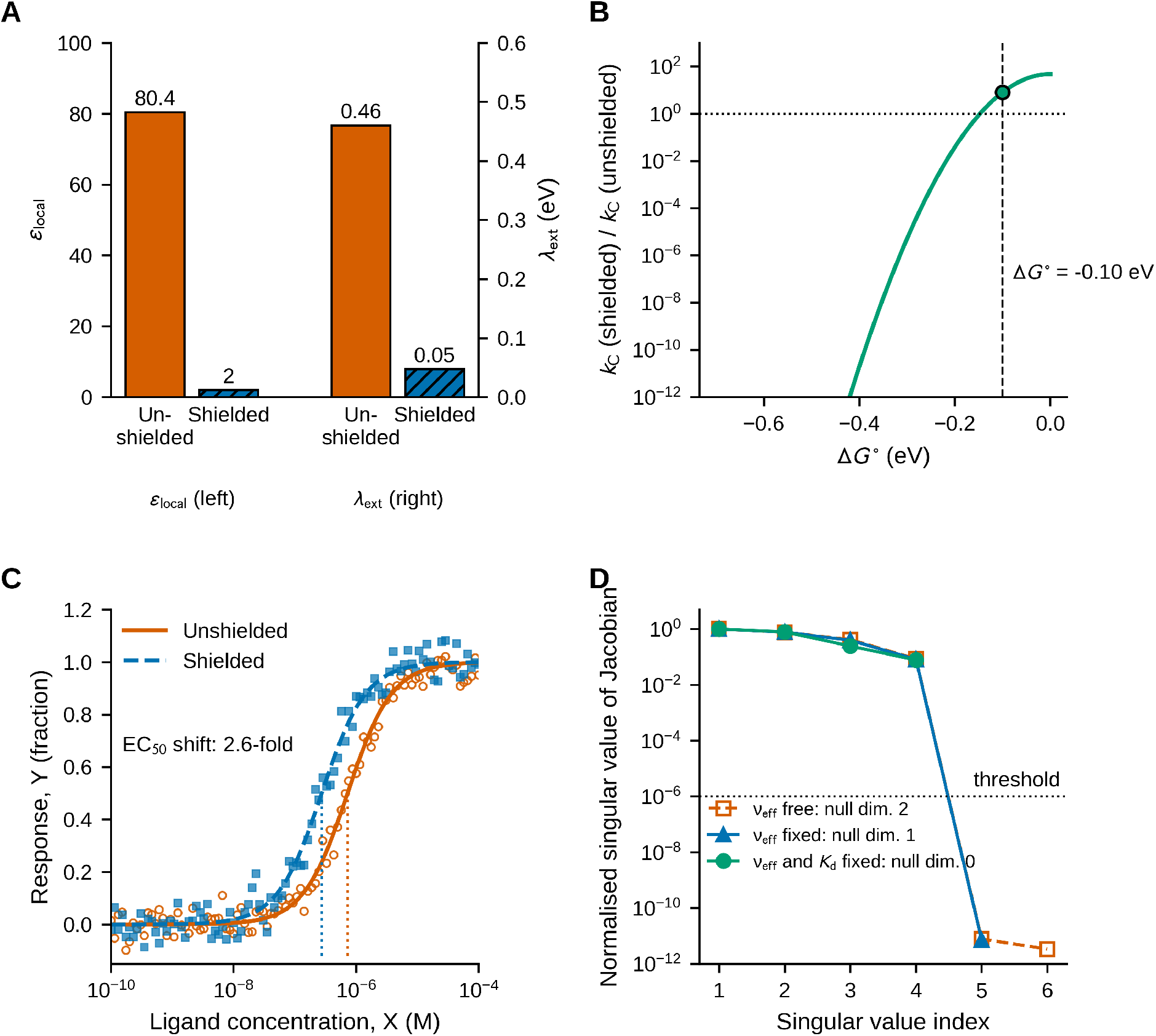
Numerical illustration of the Cluchagues-Marcus Framework. **(A)** Local dielectric constant (*ε*_local_ = 80.4 → 2.0; left axis) and external reorganization energy from Eq. 2 (*λ*_ext_ = 0.46 → 0.047 eV; right axis). **(B)** Ratio of the activation rate constants in the shielded and unshielded pocket as a function of the binding free energy Δ*G*° (log scale; dashed line, Δ*G*° = −0.10 eV used elsewhere; dotted line, ratio = 1). **(C)** Simulated dose–response curves (symbols, synthetic data with Gaussian noise, *σ* = 0.05, *N* = 100 per condition; lines, noise-free model; dotted vertical lines, EC_50_). **(D)** Normalized singular values of the Jacobian at the optimum for the three fitting configurations (representative data set, seed 42); values below 10^−6^ (dotted line) define null dimensions of the parameter space. Colors follow a color-blind-safe palette; conditions are also distinguished by line style, marker and hatching. Parameters are listed in Table 2.

#### Sensitivity analysis

The dependence of the main predictions on the assumed parameters was assessed by Monte Carlo sampling (10,000 draws; NumPy generator, seed 2026) with uniform distributions for Δ*G*° (−0.40 to −0.02 eV), *ε*_shielded_ (2–4), *ε*_opt_ (1.4–1.9), the unshielded *λ*_ext_ (0.30–0.70 eV) and *n* (0.8–2.0), and log-uniform distributions for *v*_eff_ (10^12^–10^13^ s^−1^) and *k*_off,q_ (10^10^–10^14^ s^−1^). In every draw the prefactor *G* was re-calibrated so that *λ*_ext_ at *ε*_local_ = 80.4 equalled the sampled unshielded value. The outputs were the percentage reduction in *λ*_ext_, the ratio of *k*_C_ in the shielded and unshielded pocket, and the EC_50_ fold-shift; their association with each input was quantified by Spearman rank correlation (*ρ*). All code is provided as Supplementary Code S1 and archived on Zenodo [23].

#### Real data

To test the identifiability result on measured data, napabucasin cell-viability curves for 701 cancer cell lines were analysed (eight concentrations from 0.00061 to 10 µM; file dose-data_napabucasin.csv, in the format of the DepMap portal). The data were retrieved from the Broad Institute DepMap Portal (PRISM Repurposing Secondary Screen, Public 23Q4 release) under a Creative Commons Attribution 4.0 International (CC BY 4.0) licence [24]. The response was defined as 1 − viability. Each curve was fitted with a four-parameter logistic (4PL) function (scipy.optimize.curve_fit, Trust Region Reflective algorithm, with bounds). Eq. 3 was then fitted with *k*_C_ fixed at the shielded value of Table 1 and *K*_d_ fixed at each of four assumed anchors (0.3, 1, 3 and 10 µM; free parameters Top, Bottom, *n* and *k*_off,q_). If Proposition 2 holds, the sum of squared residuals (SSE) of the anchored fits must equal that of the 4PL fit for any feasible anchor, whereas the fitted *k*_off,q_ must change. The Jacobian null dimension of the six-, five- and four-parameter configurations was also computed at the optimum of each curve. Napabucasin has been reported to act on STAT3 signalling, but its cytotoxicity is not restricted to one target, and cell viability does not measure binding to a pocket; the data were therefore used only to test fitting and identifiability, not the dielectric mechanism.

**Table 1.** Physical and kinetic quantities for the unshielded and shielded pocket (illustrative parameters of Table 2).

| **Quantity** | **Unshielded pocket** | **Shielded pocket** | **Unit** | **Ref.** |
|---|---|---|---|---|
| Local dielectric constant, *ε*<sub>local</sub> | 80.4 | 2.0 | dimensionless | [4, 9] |
| External reorganization energy, *λ*<sub>ext</sub> | 0.46 | 0.047 | eV | [3, 25] |
| Activation rate, *k*<sub>C</sub> (Δ*G*° = −0.10 eV) | 4.6 × 10<sup>11</sup> | 3.7 × 10<sup>12</sup> (8.0-fold) | s<sup>−1</sup> | [5] |
| Attempt frequency, *ν*<sub>eff</sub> | 6.46 × 10<sup>12</sup> | 6.46 × 10<sup>12</sup> (fixed) | s<sup>−1</sup> | [13] |
| Half-maximal concentration, EC<sub>50</sub> | 7.3 × 10<sup>−7</sup> | 2.8 × 10<sup>−7</sup> (2.6-fold lower) | M | — |

**Table 2.** Parameters of the simulation.

| Parameter | Value | Status |
|---|---|---|
| Temperature, *T* | 310.15 K | physiological |
| Attempt frequency, *ν*<sub>eff</sub> | 6.46 × 10<sup>12</sup> s<sup>−1</sup> | fixed *a priori* [13,14] |
| *ε*<sub>local</sub> (unshielded / shielded) | 80.4 / 2.0 | model input [4, 9] |
| Optical dielectric constant, *ε*<sub>opt</sub> | 1.8 | model input [4] |
| *λ*<sub>ext</sub> (unshielded); prefactor *G* | 0.46 eV; 0.847 eV | calibration (Section 2.1) |
| Δ*G*° | –0.10 eV | assumed |
| *K*<sub>d</sub>; *k*<sub>off,q</sub>; *n* | 1 µM; 10<sup>12</sup> s<sup>−1</sup>; 1.2 | assumed |
| Top; Bottom | 1; 0 | assumed |
| Noise *σ*; points *N*; range | 0.05; 100; 10<sup>−10</sup>–10<sup>−4</sup> M | simulation design |
| Replicates; representative seed | 200 (seeds 0–199); 42 | simulation design |

### 2.5. Software, computing environment and use of artificial intelligence

All simulations, fits and figures were produced with Python 3.12.3 (NumPy 2.4.4, SciPy 1.17.1, Matplotlib 3.10.8) on Ubuntu 24.04.4 LTS (Table 3). Random numbers were drawn with the default NumPy generator (PCG64) and the seeds given above. The Python code (Supplementary Code S1) was written and executed, and the numerical results and figures were generated, with the assistance of a large language model, Claude Sonnet 5.5 (Anthropic; model identifier claude-sonnet-5-5), accessed through the Claude chat application (claude.ai) in October 2026 using its sandboxed code-execution tool. The same model assisted with drafting the derivation and proofs (Sections 2.3 and 3.1), checking references, structuring the manuscript and language editing; the Word file was assembled with python-docx 1.2.0 and Pandoc 3.1.3. The language model is not an author and cannot assume responsibility for the work. The authors re-ran Supplementary Code S1 in Google Colab and checked the numerical results against the manuscript. No other AI tool was used in preparing the manuscript, the code or the figures.

**Table 3.**
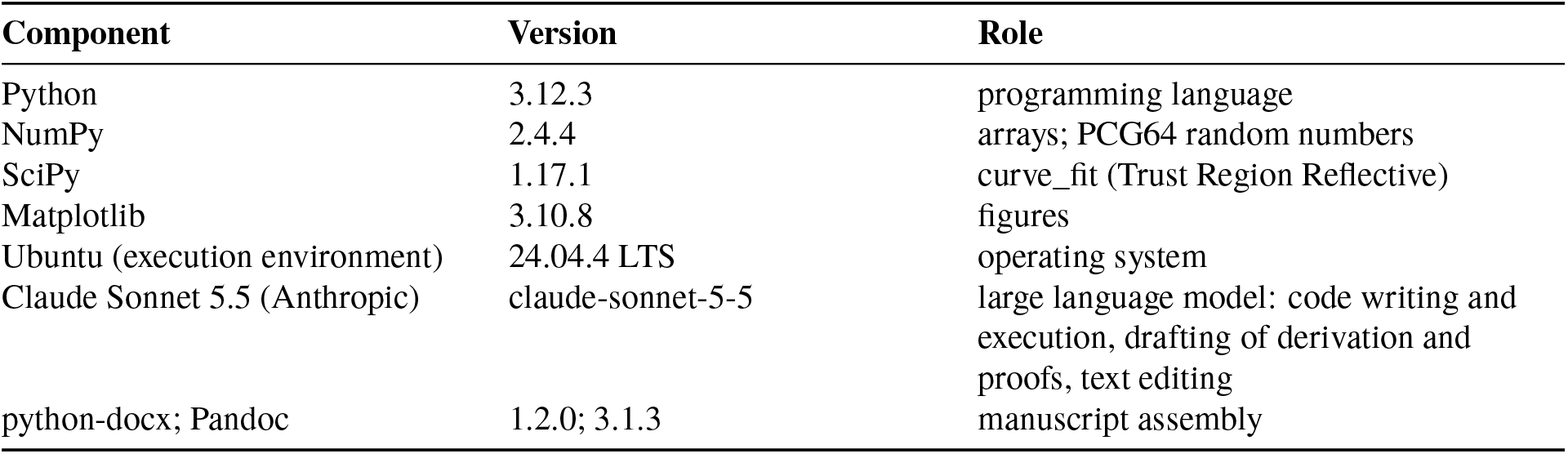
Software, language model and versions used.

## 3. Results

### 3.1. Analytical results

Throughout, *λ*_u_ and *λ*_s_ denote the external reorganization energy of the unshielded and shielded pocket, with *λ*_u_ > *λ*_s_ > 0 (which holds whenever the shielded dielectric constant lies between *ε*_opt_ and the unshielded one, by Eq. 2), and *β* = 1/ (*k*_B_*T*).

#### Proposition 1

**(rate constant)**

Let *k*_C_(λ) = *v*_eff_ exp[−(ΔG° + λ)^2^/(4λ*k*_B_*T*)]. (a) For every ΔG°, *k*_C_ ≤ *v*_eff_, with equality if and only if ΔG° = −λ. (b) *k*_C_(*λ*_s_) > *k*_C_(*λ*_u_) if and only if 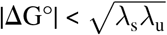; the two rates are equal when 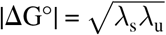, and for larger |ΔG°| shielding decreases the rate. (c) The ratio is given exactly by *k*_C_(*λ*_s_)/*k*_C_(*λ*_u_) = exp{β(*λ*_u_ − *λ*_s_)[1 − ΔG°^2^/(*λ*_u_*λ*_s_)]/4}.

*Proof*. Write *k*_C_ = *v*_eff_ exp[−*E*(λ)] with *E*(λ) = β(ΔG° + λ)^2^/(4λ) = β(ΔG°^2^/λ + 2ΔG° + λ)/4. Since *E*(λ) ≥ 0 with equality only for ΔG° = −λ, part (a) follows. For (b) and (c), *E*(*λ*_u_) − *E*(*λ*_s_) = (β/4)[ΔG°^2^(1/*λ*_u_ − 1/*λ*_s_) + (*λ*_u_ − *λ*_s_)] = (β/4)(*λ*_u_ − *λ*_s_)[1 − ΔG°^2^/(*λ*_u_*λ*_s_)]. Because *λ*_u_ > *λ*_s_, this is positive if and only if ΔG°^2^ < *λ*_u_*λ*_s_. Then *k*_C_(*λ*_s_) > *k*_C_(*λ*_u_) if and only if *E*(*λ*_s_) < *E*(*λ*_u_), which proves (b), and exponentiating gives (c).

#### Proposition 2

**(identifiability)**. Let θ = (Top, Bottom, *K*_d_, *n, k*_C_, *k*_off,q_) with Top ≠ Bottom, *n* > 0 and all rate constants positive. The response *Y*(*X*; θ) of Eq. 3 depends on θ only through (Top, Bottom, *n*, EC_50_), where EC_50_ = *K*_d_[*k*_off,q_/(*k*_C_ + *k*_off,q_)]^1/*n*^. Consequently: (a) the set of parameter vectors giving the same curve is a two-dimensional manifold, so the model has exactly two structurally non-identifiable directions; (b) if *k*_C_ is fixed (for instance from Eq. 1 with fixed *v*_eff_, ΔG° and *λ*_ext_), exactly one direction remains; (c) if *k*_C_ and *K*_d_ are both fixed, the remaining parameters (Top, Bottom, *n, k*_off,q_) are identifiable.

*Proof*. By Eq. 3, *Y* = Bottom + (Top − Bottom)/[1 + (EC_50_/*X*)^*n*^], which depends on θ only through (Top, Bottom, *n*, EC_50_). The map from the six parameters to these four quantities has rank four (the four-parameter logistic curve determines Top, Bottom, *n* and EC_50_ from its values on any interval, because Top ≠ Bottom and *n* > 0), and EC_50_ varies with each of *K*_d_, *k*_C_ and *k*_off,q_, so the level sets have dimension 6 − 4 = 2, which gives (a). Fixing *k*_C_ removes one parameter without reducing the rank, leaving 5 − 4 = 1 direction, which gives (b). For (c), with *k*_C_ and *K*_d_ fixed, *∂*EC_50_/*∂k*_off,q_ > 0 because *k*_off,q_/(*k*_C_ + *k*_off,q_) is strictly increasing in *k*_off,q_ when *k*_C_ > 0; hence *k*_off,q_ is uniquely determined by EC_50_, and (Top, Bottom, *n, k*_off,q_) is determined by the curve.

#### Proposition 3

**(consequence for potency)**. Under the assumptions of Proposition 1 and fixed *K*_d_, *k*_off,q_ and *n*, the EC_50_ in the shielded pocket is lower than in the unshielded pocket if and only if *k*_C_(*λ*_s_) > *k*_C_(*λ*_u_), that is, if and only if 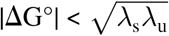.

*Proof*. EC_50_ = *K*_d_[*k*_off,q_/(*k*_C_ + *k*_off,q_)]^1/*n*^ is strictly decreasing in *k*_C_, and the result follows from Proposition 1(b).

These statements were checked numerically (Supplementary Code S1). For the calibrated values, 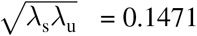 eV. In 200 random pairs of reorganization energies, the largest difference between this threshold and the numerically found crossover was 5 × 10^−6^ eV. The Jacobian of the noise-free model had rank 4 in all 500 random parameter sets for the six-, five- and four-parameter configurations, and *k*_C_/*v*_eff_ never exceeded 1.

### 3.2. Numerical illustration: dielectric collapse and rate constant

Replacing bulk water by a solvent-excluded medium lowered *ε*_local_ from 80.4 to 2.0 [4, 9]. Because Eq. 2 depends on 1/*ε*_local_, this reduced *λ*_ext_ from 0.46 eV to 0.047 eV, a decrease of 89.8% (Figure 1A, Table 1) [3, 25]. At ΔG° = −0.10 eV, *k*_C_ increased 8.0-fold (from 4.6 × 10^11^ to 3.7 × 10^12^ s^−1^) [5], in agreement with Proposition 1(c). The ratio was 30-fold at −0.05 eV, but fell below unity beyond |ΔG°| = 0.147 eV (0.038 at −0.20 eV; 5 × 10^−6^ at −0.30 eV), because the lowered *λ*_ext_ moves strongly exergonic transitions into the Marcus inverted region (Figure 1B). Acceleration is therefore conditional, and *k*_C_ cannot exceed *v*_eff_.

A comparative summary of physical and kinetic quantities is given in Table 1. In the dose–response simulation the shielded pocket shifted EC_50_ from 7.3 × 10^−7^ to 2.8 × 10^−7^ M (2.6-fold; Figure 1C), consistent with Proposition 3.

<utable>

### 3.3. Numerical illustration: parameter identifiability and optimization performance

The fits of all three configurations were equally good (median *r*^2^ = 0.988, interquartile range 0.986–0.989; median RMSE 0.048 versus the simulated noise of 0.05), so goodness of fit alone cannot distinguish between them; the *r*^2^ is limited by the noise level, and only 1% of replicates exceeded 0.992. Identifiability, however, differed (Figure 1D), as predicted by Proposition 2. With six free parameters, the Jacobian had two null dimensions. Fixing *v*_eff_ (and hence *k*_C_) reduced the null dimension to one, leaving *K*_d_ and *k*_off,q_ non-separable. Only when *K*_d_ was also fixed (for example from an independent equilibrium measurement) did the Jacobian have full rank, and *k*_off,q_ was then recovered without bias (log_10_ estimate 12.00 ± 0.06 s^−1^, mean ± SD over 200 replicates, true value 12.00). The optimizer converged within a median of 6 (interquartile range 5–6), 6 (6–7) and 7 (6–7) function evaluations for configurations (i)–(iii), respectively (excluding finite-difference Jacobian evaluations).

### 3.4. Global sensitivity analysis

Because Δ*G*°, *ε*_shielded_, *ε*_opt_, the unshielded *λ*_ext_, *v*_eff_, *k*_off,q_ and *n* are assumed rather than measured, the predictions were re-evaluated over 10,000 random parameter sets drawn from the ranges in Section 2.4 (Figure 2). Dielectric shielding lowered *λ*_ext_ by a median of 54% (5th–95th percentile, 39–80%; this is smaller than in Section 3.2 because shielded dielectric constants up to 4 were sampled). It increased *k*_C_ in 77% of the draws (median ratio 3.9; 5th–95th percentile, 0.04–27). Acceleration was almost universal for weakly exergonic binding (Δ*G*° > −0.15 eV; 99.7% of draws) and much less common for Δ*G*° ≤ −0.15 eV (65%). The rank correlation with the *k*_C_ ratio was highest for Δ*G*° (*ρ* = 0.76) and the unshielded *λ*_ext_ (*ρ* = 0.55) and weak for *ε*_shielded_, *ε*_opt_, *v*_eff_, *k*_off,q_ and *n* (|*ρ*| ≤ 0.03). The EC_50_ fold-shift (median 1.2; 5th–95th percentile, 0.37–5.6; above 2 in 26% of draws) additionally depended on *k*_off,q_ (*ρ* = −0.45), because the correction factor in Eq. 3 approaches unity when *k*_off,q_ ≫ *k*_C_. The qualitative prediction, an acceleration of *k*_C_ that depends on Δ*G*°, is therefore robust within these ranges, whereas its magnitude is not.

**Figure 2.**
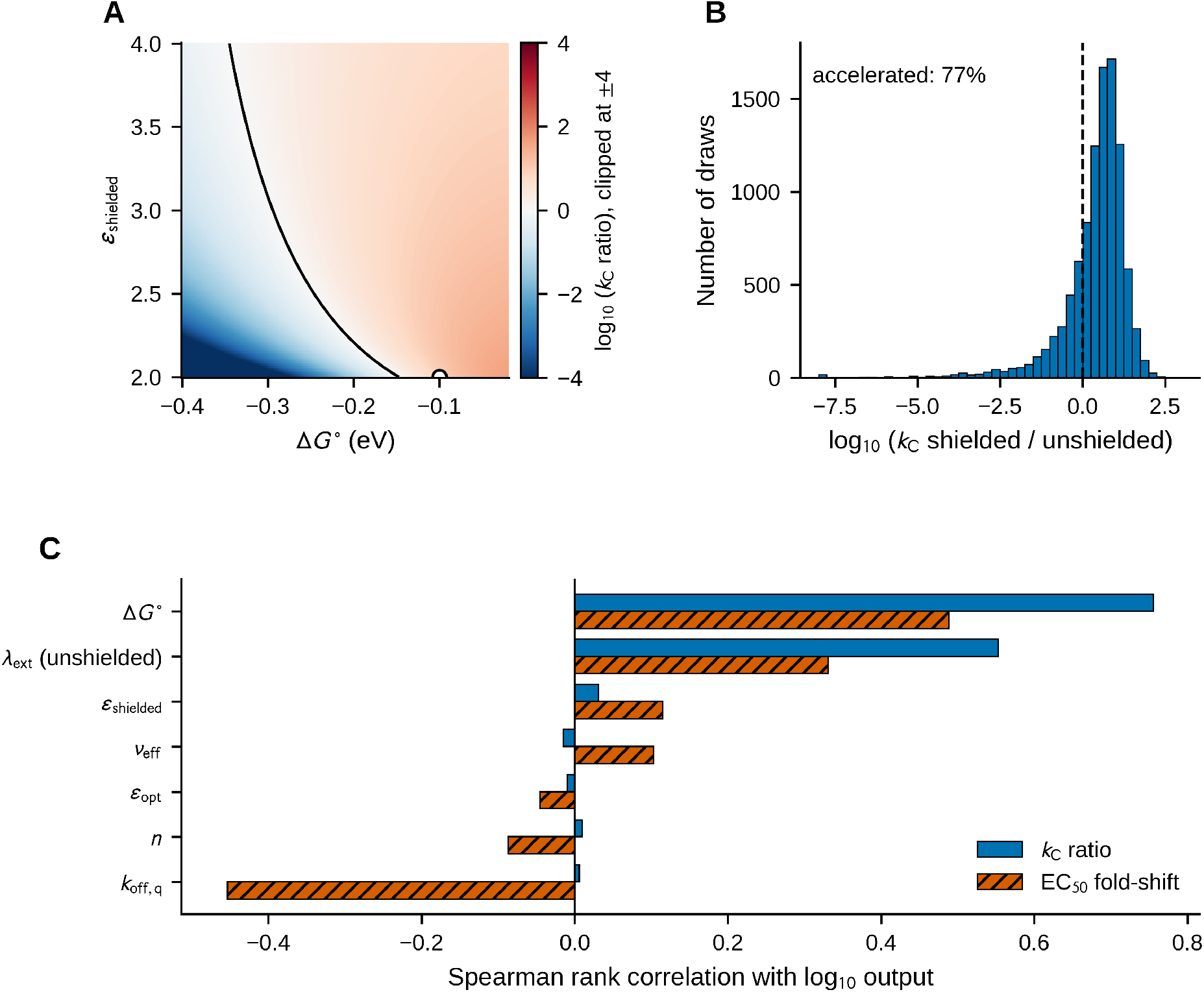
Global sensitivity analysis. **(A)** log_10_ of the *k*_C_ ratio (shielded/unshielded) as a function of Δ*G*° and *ε*_shielded_ (other parameters as in Table 2); the solid line marks a ratio of 1 and the open circle the setting used in Figure 1. **(B)** Distribution of the log_10_ *k*_C_ ratio over 10,000 random parameter sets (dashed line, ratio = 1). **(C)** Spearman rank correlations between each parameter and the log_10_ *k*_C_ ratio or EC_50_ fold-shift. Colors follow a color-blind-safe palette; bars are also distinguished by hatching.

### 3.5. Case studies: KRAS G12C and STAT3

Applying the framework to the two reference pockets (Table 4) gave the same qualitative picture. Water exclusion lowered *λ*_ext_ by 89.8% in both pockets. Because *G* scales as 1/*R* and the measured median radius is larger for STAT3 (0.83 nm) than for KRAS (0.72 nm), the STAT3 pocket had a smaller unshielded *λ*_ext_ (0.399 eV versus 0.46 eV), a smaller acceleration of *k*_C_ at Δ*G*° = −0.10 eV (3.6-fold versus 8.0-fold) and a smaller EC_50_ shift (1.9-fold versus 2.6-fold). The acceleration threshold of Proposition 1 was 0.147 eV (KRAS) and 0.128 eV (STAT3). Identifiability did not depend on the target: the null dimension was 2, 1 and 0 for the free, *v*_eff_-fixed and *v*_eff_-and-*K*_d_-fixed configurations, and fits were equally good in all three (median *r*^2^ = 0.987 for KRAS and 0.983 for STAT3). The reference dissociation constant therefore acted as the anchor that removes the last non-identifiable direction, whereas the pocket radius entered only through *G* and hence *λ*_ext_. These case studies use literature reference constants, radii measured from single crystal structures and synthetic data, and they do not constitute experimental validation.

**Table 4.** Case-study results for the two reference pockets (synthetic data; 200 replicates; parameters of Table 2).

| Quantity | KRAS G12C (6UT0) | STAT3 SH2 (6NJS) |
|---|---|---|
| Pocket radius *R*; reference constant | 0.72 nm; 1.43 µM (*K*<sub>I</sub>, adagrasib) | 0.83 nm; 11 nM (*K*<sub>i</sub>, SD-36) |
| Prefactor *G* | 0.847 eV | 0.735 eV |
| λ<sub>ext</sub>: unshielded → shielded | 0.46 → 0.047 eV | 0.399 → 0.041 eV |
| Reduction of λ<sub>ext</sub> | 89.8% | 89.8% |
| *k*<sub>C</sub> ratio at Δ*G*° = –0.10 eV | 8.0-fold | 3.6-fold |
| EC<sub>50</sub> fold-shift | 2.6 | 1.9 |
| Threshold $\sqrt{\lambda_{s}\lambda_{u}}$ | 0.147 eV | 0.128 eV |
| Null dimension: free / *ν*<sub>eff</sub> fixed / *ν*<sub>eff</sub> and *K*<sub>d</sub> fixed | 2 / 1 / 0 | 2 / 1 / 0 |
| Median *r*<sup>2</sup> | 0.987 | 0.983 |
| *k*<sub>off,q</sub> recovered with *ν*<sub>eff</sub> and *K*<sub>d</sub> fixed (log<sub>10</sub>, mean ± SD) | 12.00 ± 0.06 | 11.99 ± 0.05 |

### 3.6. Real-data test of the identifiability result

The 4PL fits described the 701 napabucasin curves with a median *r*^2^ of 0.91 (interquartile range 0.85– 0.95), lower than for the simulated data because real curves are noisier. The fitted EC_50_ agreed with the EC_50_ reported in the data file (Spearman *ρ* = 0.86; median absolute difference 0.08 log_10_ units; Figure 3B). For every curve, the Jacobian null dimension of the full model was 2, 1 and 0 for the six-, five- and four-parameter configurations, as in the simulations. Anchoring *K*_d_ at 0.3, 1, 3 and 10 µM (feasible for 258, 612, 687 and 693 curves, respectively, because EC_50_ cannot exceed *K*_d_ in Eq. 3) left the SSE unchanged relative to the 4PL fit (median ratio 1.0000; 95th percentile ≤ 1 + 10^−11^), while the median fitted log_10_ *k*_off,q_ changed from 12.6 to 9.1 (Figure 3C). Real viability data therefore cannot determine *K*_d_ or *k*_off,q_ separately, which is the non-identifiability stated in Proposition 2. These curves serve as a numerical stress test of fitting and identifiability on real noisy data, not as biological validation, and they do not test the dielectric mechanism.

**Figure 3.**
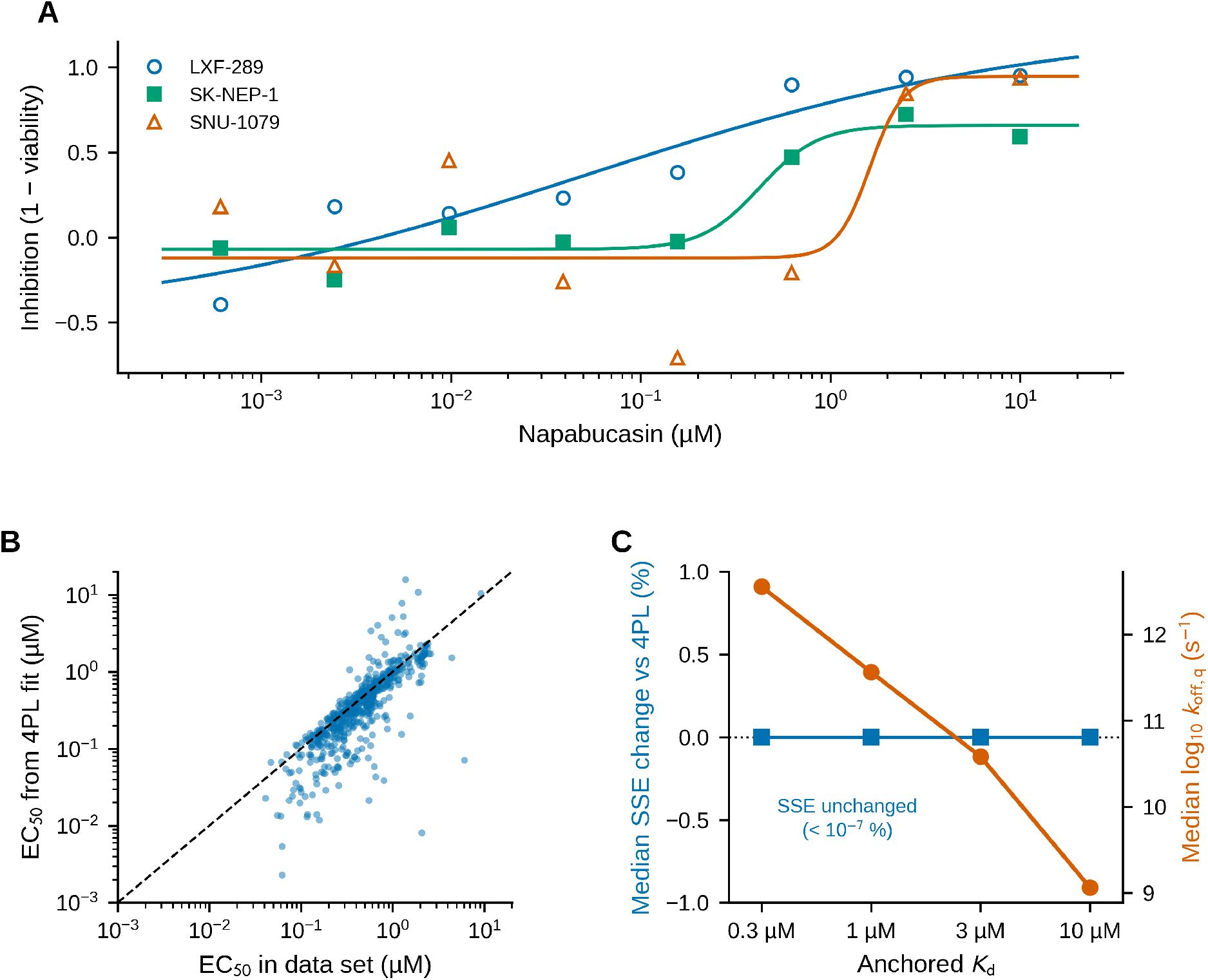
Real-data test (napabucasin, 701 cancer cell lines). **(A)** Example curves (5th, 50th and 95th percentile of the fitted EC_50_) with four-parameter logistic fits. **(B)** EC_50_ from the 4PL fits against the EC_50_ reported in the data file (dashed line, identity). **(C)** Median change of the sum of squared residuals relative to the 4PL fit (blue, left axis) and median fitted log_10_ *k*_off,q_ (orange, right axis) when *K*_d_ is anchored at four values. The fit quality does not change while *k*_off,q_ varies by 3.5 log_10_ units. Colors follow a color-blind-safe palette.

## 4. Discussion

The mathematical results are of three kinds. First, Proposition 1 gives the exact boundary 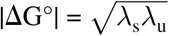 between acceleration and deceleration of the Marcus rate constant when the reorganization energy is lowered; the intuition is that a lower reorganization energy raises the rate only while the transition is not already beyond the barrierless point. Second, Proposition 2 shows that the kinetically corrected Hill model carries only four identifiable combinations of its six parameters, so no fitting algorithm can recover *K*_d_, *k*_C_ and *k*_off,q_ separately from dose–response data alone, and fixing a physical constant such as *v*_eff_ reduces but does not remove the ambiguity [6,7, 13]. Third, the sensitivity analysis shows that the numerical size of the effects depends mainly on Δ*G*° and the unshielded *λ*_ext_.

The case studies show how a measured dissociation constant and a pocket geometry would enter the framework in practice: the former anchors the potency parameter and the latter scales the reorganization energy. A numerical stress test on real napabucasin viability curves was consistent with the identifiability result: fit quality was insensitive to the dissociation-constant anchor while the fitted off-rate changed by orders of magnitude. Proposition 2 concerns structural identifiability, a property of the equations evaluated here on noise-free derivatives; practical identifiability, the ability to extract parameters from noisy and sparse data, is a separate property. With eight concentrations and noisy viability curves (median *r*^2^ = 0.91), even the four identifiable quantities carry substantial uncertainty, and parameter confidence intervals were not evaluated. These results concern the model, not a biological mechanism. Whether dielectric shielding changes potency in a real pocket would require experimental or structure-based evidence that is not provided here.

### 4.1. Limitations

First, the data used to illustrate the framework are synthetic and generated from the same model that is then fitted, so recovery of parameters demonstrates mathematical properties of the formulation, not biological validity [21, 26]. The real data (Section 3.6) are cell-viability curves of a compound whose action is not restricted to a single pocket, so they test the fitting and identifiability but not the dielectric mechanism or binding kinetics. Second, Marcus theory was derived for electron transfer, and its extension to ligand– pocket transitions is an assumption that has not been validated experimentally. Moreover, Eq. 3 treats *k*_C_ and *k*_off,q_ as independent, which is justified only for an open (driven) scheme such as the steady-state scheme of Section 2.3; in a closed equilibrium scheme, detailed balance would fix their ratio at exp (−Δ*G*°/*k*_B_*T*), and the equilibrium EC_50_ would then not change with *λ*_ext_ alone, only through a change in Δ*G*°. Third, Δ*G*°, *k*_off,q_, *n* and the prefactor *G* were assumed or calibrated. In the case studies, the pocket radii are single-structure geometric estimates (median ligand-to-pocket distance) rather than cavity radii, and the KRAS constant is a covalent inhibition constant (*K*_I_) rather than a reversible dissociation constant. Each constant belongs to the ligand present in the corresponding structure (adagrasib in 6UT0 [16]; SD-36 in 6NJS [19]), but the two constants come from different assays (mass spectrometry and fluorescence polarization). Their influence was explored in a sensitivity analysis (Section 3.4), but they remain unconstrained by experimental data. In that analysis Δ*G*° and *λ*_ext_ were sampled independently, whereas in real pockets mutations or chemical substitutions that change the local dielectric boundary also change the electrostatic contribution to binding, so Δ*G*° and *λ*_ext_ are covariates. Fourth, the value *k*_off,q_ = 10^12^ s^−1^ was chosen so that *k*_off,q_ is comparable with *k*_C_, because otherwise the correction factor in Eq. 3 is trivially close to unity. It should be read as the microscopic rate at which the engaged state is removed (Section 2.3), that is, a pocket relaxation rate (for example structural relaxation or re-entry of water) on the timescale of molecular vibrations, not as the experimentally measured dissociation rate. Real ligands typically show dissociation rates of about 10^−6^–10^3^ s^−1^, and equating *k*_off,q_ with the measured rate would, for *K*_d_ = 1 μM, imply an association rate of about 10^18^ M^−1^ s^−1^, far above the diffusion limit (about 10^9^–10^10^ M^−1^ s^−1^). The physical meaning of *k*_off,q_ and its relation to the macroscopic dissociation rate therefore require justification in future work. Fifth, continuum dielectric models become unreliable at sub-nanometer length scales, where hydrogen-bonding networks, conformational flexibility and entropic penalties dominate [20], and the existence, stability, delivery and toxicity of sub-nanometer metal vectors are not addressed. The continuum treatment is a macro-scale abstraction, and molecular dynamics (MD) simulations with explicit solvent are required to evaluate the microscopic solvent configurations inside tight cavities. Future work should test the framework against real dose–response data and kinetic measurements, for example by surface plasmon resonance (SPR), and against structure-based calculations such as molecular dynamics (MD) simulations [6, 27].

## 5. Conclusions

For a Marcus rate constant entering a kinetically corrected Hill model, lowering the reorganization energy increases the rate if and only if 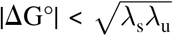, and the rate never exceeds the attempt frequency. The model has exactly two non-identifiable parameter directions, which are removed by fixing the attempt-frequency-derived rate constant and anchoring the dissociation constant, as shown for KRAS G12C and STAT3 reference pockets. Numerical illustrations and a global sensitivity analysis are consistent with these results. A numerical stress test on real napabucasin viability curves is consistent with the identifiability result but is not biological validation; the dielectric mechanism has not been tested against biological data and the framework remains hypothesis-generating.

## Acknowledgements

The authors acknowledge the use of Claude Sonnet 5.5 (Anthropic) for code generation, numerical analysis support and language editing, as described in Section 2.5.

## Statements and Declarations

### Ethical approval

Not applicable. This study involved purely computational modeling and did not use human participants, patient data or animal models.

### Data and code availability

All code needed to regenerate the synthetic data sets, fits, statistics, propositions checks and figures (Supplementary Code S1; Python, fixed random seeds) is openly available at https://github.com/cluchagues23-rgb/Cluchagues-Marcus-framework and archived on Zenodo (version v1; https://doi.org/10.5281/zenodo.23653778; all versions: https://doi.org/10.5281/zenodo.23653777) [23]. The code is released under the MIT licence (see the repository), and the Zenodo record is licensed under CC BY 4.0. The napabucasin data file used in Section 3.6 (DepMap Public 23Q4, CC BY 4.0 [24]) is included in the repository. No other data were generated.

### Intellectual property

The authors have not filed and do not intend to file a patent application on the methods described in this article. The code is released under the MIT licence.

### Author contributions (CRediT)

L. M. Cluchagues Abam: Conceptualization, Methodology, Writing – original draft, Project administration. I. H. Mfabo Kameni: Validation, Formal analysis (review of the code and results), Writing – review & editing. F. Eya’ane Meva: Supervision, Writing – review & editing. All authors approved the final manuscript and agree to be accountable for the work.

### Use of artificial intelligence

A large language model (Claude Sonnet 5.5, Anthropic) was used for code generation and execution, numerical analysis, drafting of the derivation and proofs, figure generation, reference checking, structuring and language editing, as described in Section 2.5 and Table 3. No AI tool is listed as an author.

## Abbreviations

MD: molecular dynamics
RMSE: root-mean-square error
SPR: surface plasmon resonance
TRR: Trust Region Reflective.

